# Automated Ventricle Assessment via Three-dimensional Anatomical Reconstruction (AVA-TAR): a computational toolkit for autonomous lateral ventricle assessment in preclinical hydrocephalus models

**DOI:** 10.64898/2026.02.02.703412

**Authors:** Sundeep Chakladar, Shelei Pan, Owen Limbrick, Maneesha Pandey, Grace L. Halupnik, Annie Zhao, Moe R. Mahjoub, James D Quirk, Arash Nazeri, Jennifer M. Strahle

## Abstract

**Introduction:** Current workflows for studying hydrocephalus in rodent models rely on manual segmentation or qualitative assessment of ventricular size on small animal magnetic resonance imaging, which are both inefficient and prone to variability. Atlas-based methods enable more streamlined segmentation, but their analysis is limited to morphologically normal samples.

**Objective:** This study aimed to develop and internally validate a deep learning model that performs automated segmentation of lateral ventricles in rodent brain MRIs, allowing for 3D ventricle reconstruction, morphological analysis, and ventriculomegaly detection.

**Methods:** Four U-Net++ neural networks, each with different encoder backbones, were trained using 307 rodent brain MRIs (262 rats, 45 mice), each with manually segmented lateral ventricles serving as the ground truth. Model performance was evaluated using the Dice coefficient, intersection over union (IoU), and Hausdorff index. The most optimal model was evaluated further for its ability to quantify ventricle volume, convexity, surface area, and symmetry.

**Results:** The U-Net++ model with an EfficientNet-B1 encoder achieved high accuracy (Dice: 0.823 ± 0.136; IoU: 0.721 ± 0.85). Further assessment of its morphological predictions found strong correlations with manual measurements of ventricular morphology, with Pearson and interclass correlation coefficients exceeding 0.96 across all metrics. The full validated pipeline was packaged into a publicly available application, hosted at https://ava-tar.org.

**Conclusion:** This study introduces a deep learning tool for automated segmentation and morphological analysis of lateral ventricles in rodent MRIs. The tool’s efficiency and accuracy in quantifying ventricle morphology offers significant utility in preclinical hydrocephalus research with potential future application in the clinical setting.

## INTRODUCTION

Hydrocephalus comprises a multifactorial buildup of cerebrospinal fluid (CSF) in the ventricles of the brain resulting from heterogenous etiologies with unknown pathophysiology. Hydrocephalus presents a significant neurosurgical burden, highlighting the importance of ongoing research into its mechanisms^1,2^. Preclinical hydrocephalus research encompasses both genetic mouse models of congenital hydrocephalus and surgical rodent models of hydrocephalus including post-hemorrhagic hydrocephalus. Models such as these have led to advances in understanding hydrocephalus as a neurodevelopmental disorder^3,4^, uncovering the role of inflammatory activation in post-hemorrhagic ventricular dilation ^5–12^, and exploring the role of prophylactic iron chelation^13,14^.

A common metric for assessing disease severity, progression, and treatment response in these models is longitudinal monitoring of the CSF spaces on small animal MRI and quantification via segmentation-based analyses^15^. Accurate measurement of ventricle volumes from these images is therefore integral to the analysis of these data. Despite the importance of ventricle quantification in rodent brain MRIs, current methods for doing so have significant limitations.

Commonly used approaches include qualitative visual assessment, manual segmentation, and automated atlas-based segmentation^16–19^. Visual approximation is efficient but is inherently subjective, imprecise, and lacks the quantitative rigor needed for robust statistical analysis. Manual segmentation is considered the gold-standard for accuracy, but it is inefficient, requires significant technical expertise, and is prone to inter-observer variability ^20^. More recently, atlas-based segmentation has emerged as an automated approach aimed at overcoming these issues of inefficiency and variability. This technique works by registering a standard digital brain atlas to the subject’s MRI scan. Once aligned, the anatomic labels from the atlas are transferred to the subject’s scan, thereby generating an automated segmentation ^21,22^. While atlas-based segmentation offers a more efficient and standardized alternative, its utility is limited in hydrocephalus research. The severe ventriculomegaly seen in hydrocephalus often results in distortion of the ventricular system and surrounding parenchymal structures, resulting in an MRI that looks significantly different than the standard MRI used as the atlas reference ^23^.

Convolutional neural networks (CNNs) are a class of deep learning algorithms that offer a solution to overcome these challenges. Unlike atlas-based methods, CNNs can be trained on diverse datasets to learn the features of specific anatomical structures. This allows for segmentation to be performed with high accuracy, even in the presence of significant anatomic variation across inputs ^24,25^. Consequently, CNNs can achieve the efficiency and standardization of automated methods while retaining the flexibility and precision needed to handle anatomically distorted brains, effectively combining the strengths of previous approaches.

This project aimed to 1) develop a fully automated, CNN-based pipeline for the segmentation, 3D reconstruction, and morphological quantification of lateral ventricles in rodent brain MRIs with varying levels of ventriculomegaly, 2) validate the pipeline’s segmentation and quantification accuracy, and 3) integrate the pipeline into an openly available, user-friendly web application.

## METHODS

### Animals

All animal experiments were approved by the Institutional Animal Care and Use Committee at Washington University in St. Louis (protocols #22-0416, #24-0259). Sprague Dawley Rats (crl:SD 400, Charles River Laboratories, Wilmington, MA) were used in all experiments involving rats. Rat pups were housed with their moms in a 12□h light–dark cycle (6 a.m. lights on/6 p.m. lights off) in a temperature and humidity-controlled room. Water and food were provided ad libitum for the mom. For mice experiments, generation of FoxJ1-CreERT2-Cep120F/F animals and littermate controls were done by crossing Cep120F/F mice 26–28 with the Tamoxifen-inducible Foxj1-CreERT2-GFP 29 transgenic strain. To induce Cre expression, intraperitoneal injections of Tamoxifen were administered to mice at either P0 (50 µg/kg daily for 3 days) or 8 weeks (100 mg/kg daily for 5 days).

### Intraventricular hemorrhage

To induce post-hemorrhagic hydrocephalus (PHH), we adapted our previously reported neonatal intraventricular hemorrhage (IVH) model ^26–28^. Postnatal day (P)1, 4, and 15 rats were first anesthetized with inhaled isoflurane (3% induction, 2% maintenance), and a 1 cm midline skin incision was made to expose the bregma. Rats were injected with 20 µL of 150 mg/mL hemoglobin (HB) at an 8 µL/minute rate with the following stereotaxic coordinates relative to bregma: 1.0 mm lateral, 0.0 mm anterior, and 2.0 mm deep at P1; 1.5 mm lateral, 0.0 mm anterior, and 2.0 mm deep at P4; and 1.5 mm lateral, 0.0 mm anterior, and 2.5 mm deep at P15. Age-matched rats were injected with an equivalent volume of artificial CSF to induce the control condition. The needle was left in place for 2 minutes after the end of injections to prevent backflow. Rats were sutured, allowed to recover from anesthesia, and returned to their moms.

### Magnetic resonance imaging acquisition

For rat experiments, T2-weighted MRIs were acquired at various timepoints after PHH induction ranging from 6 hours to 8 weeks to monitor for ventriculomegaly and posthemorrhagic hydrocephalus. For mice, T2-weighted MRIs were acquired at postnatal day 31. All MR images were acquired at the Washington University Small Animal Magnetic Resonance Facility on a Bruker Biospec 9.4-T MRI scanner with either a 4-channel array mouse brain CryoProbe, 4-channel mouse brain surface coil, or 4-channel rat brain surface coil. Rodents were anesthetized with isoflurane (3% induction, 2% maintenance) throughout image acquisition, and body temperature was maintained with a circulating warm waterbed. T2-weighted fast spin-echo sequences with the following parameters were used to obtain images of the entire brain: echo time 66 ms; repetition time 4938 ms; averages 2; echo spacing 16.5 ms; rare factor 8; slice thickness 0.5; FOV 15 mm x 15 mm (<P10), 18 x 18 mm (P14-P28), or 20 x 20 mm (P28+); Image size 192 x 192 or 128 x 128. Image resolution ranged from 78 x 78 x 500 µm3 and 195 x 195 x 500 µm3. 30 to 56 axial slices were acquired, covering the entire brain from olfactory bulb to cerebellum. MR images were exported as NIFTI files.

### Manual segmentation

NIFTI files were opened individually in ITK-SNAP (https://www.itksnap.org/pmwiki/pmwiki.php) for manual segmentation. The paintbrush tool was used to outline the ventricle, cavum septum pellucidum, posterior white matter edema, and any other pathological CSF space that resulted directly from ventricular expansion. All measurements were performed by an experienced observer blinded to the experiment groups. The volume of the segmentation was calculated and reported by ITK-SNAP in mm^3^.

### Lateral Ventricle Segmentation Model Development

The aggregated T2-weighted MRIs and their corresponding lateral ventricle segmentation masks were randomly divided into training, validation, and testing datasets, with a split ratio of 70:10:20. The training set was used to train four custom U-Net++ machine learning models, each with a unique encoder ^29^. The ResNet34, ResNeXt-50 (32×4d), EfficientNet-B1, and DenseNet-121 encoders were selected as candidates due to their previously demonstrated successes in similar medical imaging segmentation tasks ^30–37^. Each model was trained for 20 epochs with a learning rate of 0.0001 and batch size of 4. The comprehensive set of parameters used for model development is listed in Supplementary Table 1. The validation dataset was used to assess performance during model training, with validation loss guiding learning rate scheduling. Training loss and validation loss were monitored across epochs to ensure a plateau in model performance was reached prior to termination of learning (Supplementary Figure 1).

### Selection of Optimal Encoder

Each of the four lateral ventricle segmentation models was evaluated on the held-out testing set of MRIs. Accuracy was determined by comparing predicted segmentation masks with the manually generated masks. Pixel-level performance was quantified using the Dice score and intersection over union (IoU), both of which range from 0 to 1, with higher values indicating greater pixel-level accuracy ^38^. To assess species-specific and overall performance, Dice score and IoU were calculated separately for the rat and mouse subsets of the testing data, as well as for the full testing dataset containing both species.

To further assess encoder performance beyond pixel-level overlap, the three-dimensional accuracy of their output segmentations was also evaluated. The predicted and manual segmentation masks for each MRI in the testing set were first reconstructed into 3D volumes. The 3D Hausdorff distance was then calculated to quantify the maximum distance between the surfaces of the predicted and corresponding manual volumes, with lower values indicating a more accurate reconstruction ^39^. Consistent with the pixel-level analysis, this comparison was performed separately for the rat and mouse subsets, as well as for the entire testing dataset. To complement the quantitative metrics, a qualitative visual assessment was performed to compare anatomic similarity across the encoder outputs. Three representative rat MRIs with varying ventricle volumes were selected from the testing dataset. 3D reconstructions were generated for each of these samples using both their manual segmentation masks as well as the corresponding predictions from each of the four encoders. The resulting five volumes for each sample were visually compared to identify any variance, inaccuracies, artifacts, or systemic differences in segmentation quality.

### Intracranial Space Segmentation Model Development

To enable the normalization of lateral ventricle volumes, a second U-Net++ model was developed to segment the total intracranial space. Manual segmentation masks for the intracranial space were created for a subset of the MRI dataset. This model utilized the EfficientNet-B1 encoder, which was identified as the most optimal encoder in the lateral ventricle segmentation task. The model was trained and validated following an identical procedure to the ventricle models, using the same 70:10:20 data split and hyperparameters, ensuring a standardized development approach across both segmentation tasks.

### Accuracy of Morphological Quantification

Following the identification of the U-Net++ model with an EfficientNet-B1 encoder as the optimal architecture, we performed a secondary analysis to evaluate its ability to accurately quantify clinically important morphological features. Five metrics were calculated from the 3D reconstructions of both the manually segmented masks (ground truth metrics) and their corresponding model-predicted masks (predicted metrics). These metrics included: 1. Volume: The total volume of the lateral ventricles (in mm^3^), 2. Normalized Volume: The ratio of the lateral ventricle volume to the total intracranial volume, 3. Surface Area: The total surface area of the 3D ventricle mesh (in mm^2^), 4. Symmetry: The volumetric similarity of the ventricles across the sagittal plane, calculated as the Dice score between the left and right hemispheres, 5. Convexity: A ratio of the ventricle volume to the volume of its 3D convex hull (Figure 1).

**Figure 1.**
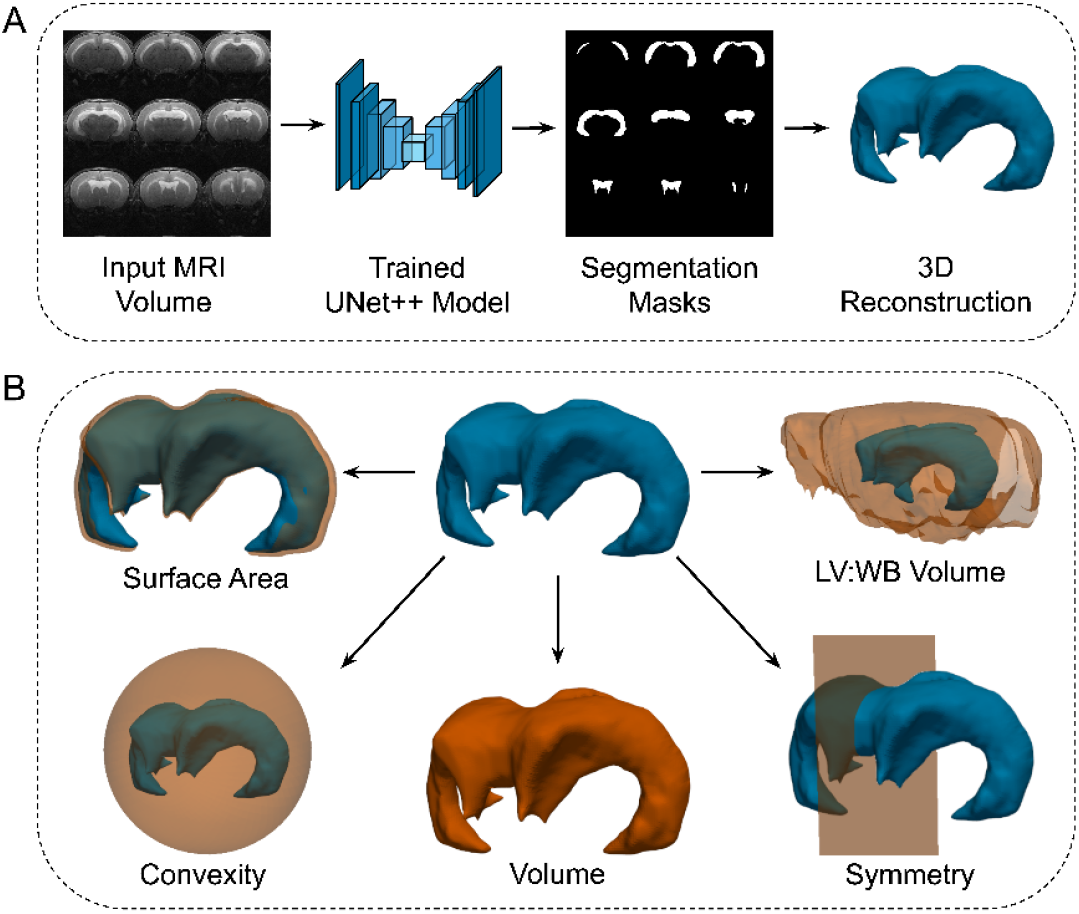
Overview of the automated ventricle analysis pipeline, including **A)** segmentation with 3D reconstruction and **B)** morphology assessment. LV:WB Volume = fraction of whole brain volume occupied by lateral ventricles.

The agreement between the predicted metrics and ground truth metrics was assessed using interclass correlation coefficient, mean absolute error, Pearson correlation coefficient, and Bland-Altman analysis. Metrics were assessed separately for rats and for mice.

### Web-Based Implementation and Public Availability

To enhance accessibility and reproducibility, the validated segmentation and analysis pipeline was packaged into a publicly available application, hosted at https://ava-tar.org. The platform provides a user-friendly, no-code interface that allows researchers to upload T2-weighted rat or mouse MRIs in the NIfTI (.nii or .nii.gz) format. The website executes automated lateral ventricle segmentation, displaying results for the user. Users can then view the five calculated morphology metrics, interact with a 3D reconstruction of the ventricles, and download the resulting segmentation mask for further offline analysis. For higher-throughput needs, the website features a batch processing mode to streamline the processing of multiple files. The batch processing mode can also be used to download the total intracranial volume segmentation mask.

### Software

Python v3.12.5 (Python Software Foundation, Wilmington, Delaware) was used for model construction, morphology quantification, performance assessment, and website backend development. All models were trained in Google Colab using an A100 GPU. RStudio v4.3.2 (RStudio, Boston, MA) was used for model performance assessment.

## RESULTS

### Image Datasets

To develop and validate the lateral ventricle (LV) segmentation model, 438 T2-weighted MRIs (376 rats, 62 mice) and their corresponding manual segmentation masks were compiled for a total of 17,583 individual slices (Figure 2). Data were randomly split into training (307 scans, 12,313 total slices), validation (43 scans, 1,730), and test (88 scans, 3,540 total slices) sets (Figure 2).

**Figure 2.**
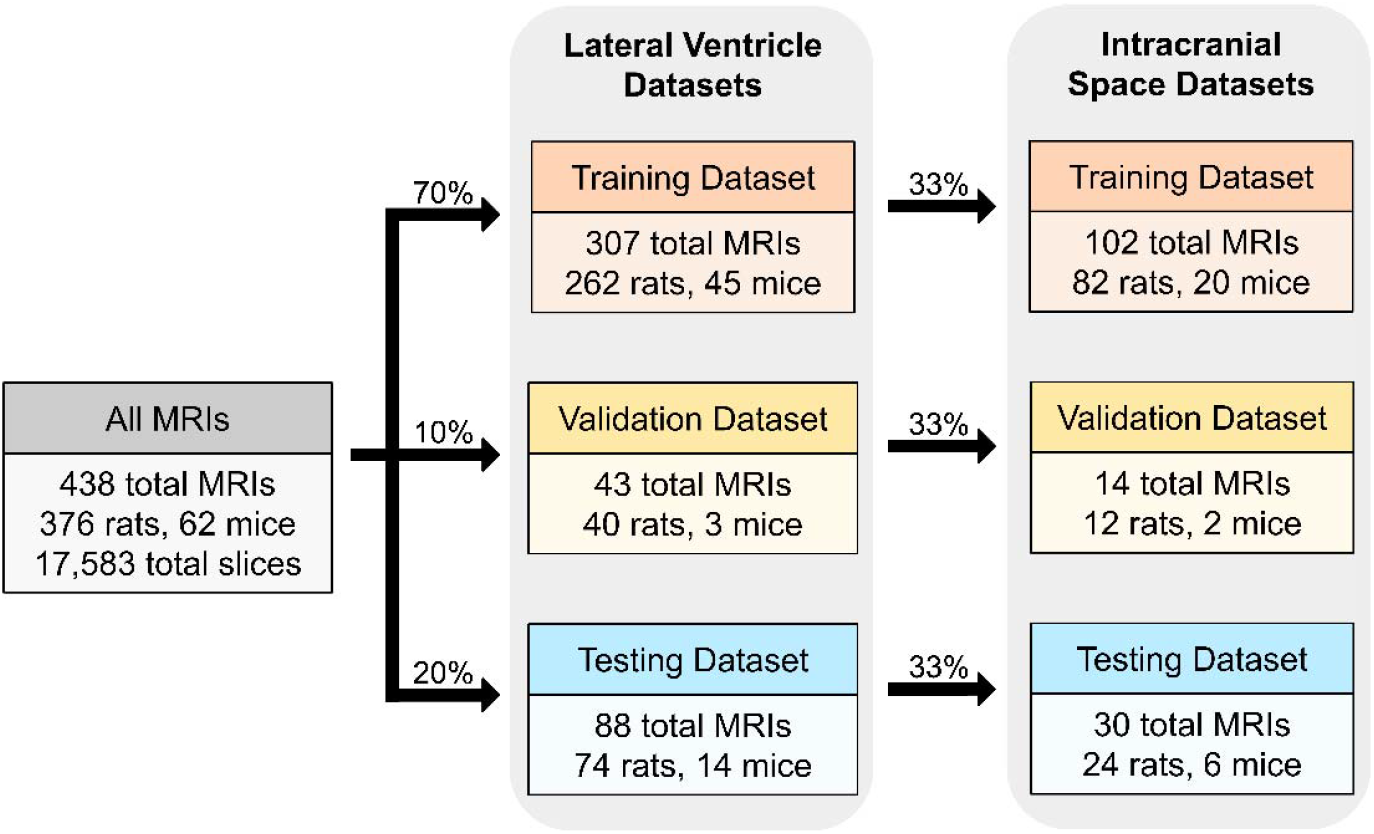
Composition of the training, validation, and testing datasets for the lateral ventricle and intracranial space segmentation models.

To incorporate downstream ventricle-to-brain (VBR) ratio analysis of the LV segmentation model output, we also developed a total intracranial volume (ICV) segmentation model. 33% of the MRIs used for the LV segmentation model were randomly selected and their corresponding IVC manual segmentation masks were compiled for a total of 5269 individual slices. Data were split into training (102 scans, 4,103 total slices), validation (14 scans, 562 total slices), and test (30 scans, 1204 total slices) sets (Figure 2).

Importantly, our diverse training dataset allows AVA-TAR to generalize to a wide range of ventricle volumes, morphologies, and rodent ages. Lateral ventricle volumes ranged from 0.6 to 2756.0 mm^3^, with a mean of 133.7 mm^3^ (SD 293.6 mm^3^) and median 16.7 mm^3^ (IQR: 7.1–94.5 mm^3^). Total intracranial volumes ranged from 28.8 to 8705.1 mm^3^, with mean of 1483.4 mm^3^ (+/-1101.7 mm^3^) and median 1186.4 mm^3^ (25th–75th percentile: 694.8–2216.9 mm^3^).

### Selection of Optimal Encoder

We used UNet++ as the base architecture for our model because of its proven effectiveness and widespread adoption in medical image segmentation ^29,40^. To identify the most suitable encoder for our semantic segmentation task, we evaluated four encoders with unique methods of feature extraction: ResNet-34, ResneXt50-32×4d, Efficientnet-b1, Densenet-121.

When evaluated on the testing dataset, the four candidate encoders achieved Dice scores ranging from 0.822 to 0.827 and IoU values ranging from 0.719 to 0.724, with no significant differences observed between encoders (Table 1).

**Table 1:** Performance Metrics for Each U-Net++ Variant.

| Encoder Backbone | Overall |  | Rats |  | Mice |  |
| --- | --- | --- | --- | --- | --- | --- |
|  | Dice Score | IoU | Dice Score | IoU | Dice Score | IoU |
| ResNet-34 | 0.822 ± 0.137 | 0.719 ± 0.188 | 0.817 ± 0.137 | 0.712 ± 0.191 | 0.849 ± 0.142 | 0.756 ± 0.182 |
| ResNeXt50-32x4d | 0.827 ± 0.129 | 0.724 ± 0.180 | 0.822 ± 0.129 | 0.717 ± 0.183 | 0.853 ± 0.133 | 0.762 ± 0.174 |
| Efficientnet-b1 | 0.823 ± 0.136 | 0.721 ± 0.185 | 0.817 ± 0.140 | 0.712 ± 0.191 | 0.858 ± 0.116 | 0.766 ± 0.159 |
| Densenet-121 | 0.825 ± 0.134 | 0.724 ± 0.184 | 0.819 ± 0.138 | 0.715 ± 0.189 | 0.860 ± 0.120 | 0.770 ± 0.163 |

The same set of testing MRIs were used to assess all models. IoU: intersection over union

Comparison of 3D Hausdorff distance across the four encoders revealed no significant differences in performance. Consistent with the pixel-level analysis, model outputs were largely comparable between encoders. Hausdorff distance measurements indicated generally stronger performance on rat samples compared to mouse samples across all encoders (Figure 3).

**Figure 3.**
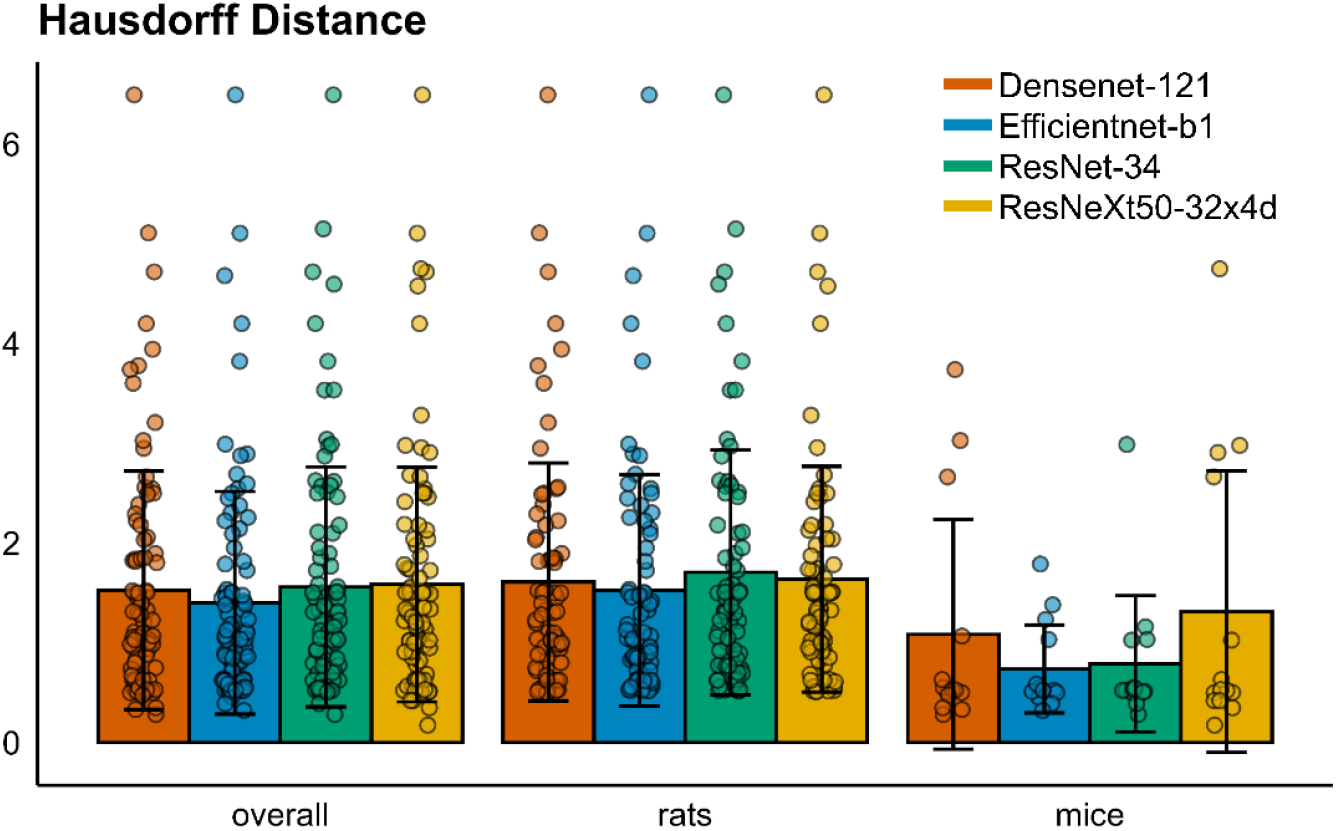
Comparison of Hausdorff distances as quantified by the four UNet++ variants.

Qualitative comparison of the 3D reconstructions produced by each encoder similarly showed no apparent differences in segmentation quality. Across all three representative samples with varying ventricle sizes, predicted reconstructions were visually consistent and closely matched the corresponding manual segmentations. No encoder appeared to exhibit systematic errors (Figure 4).

**Figure 4.**
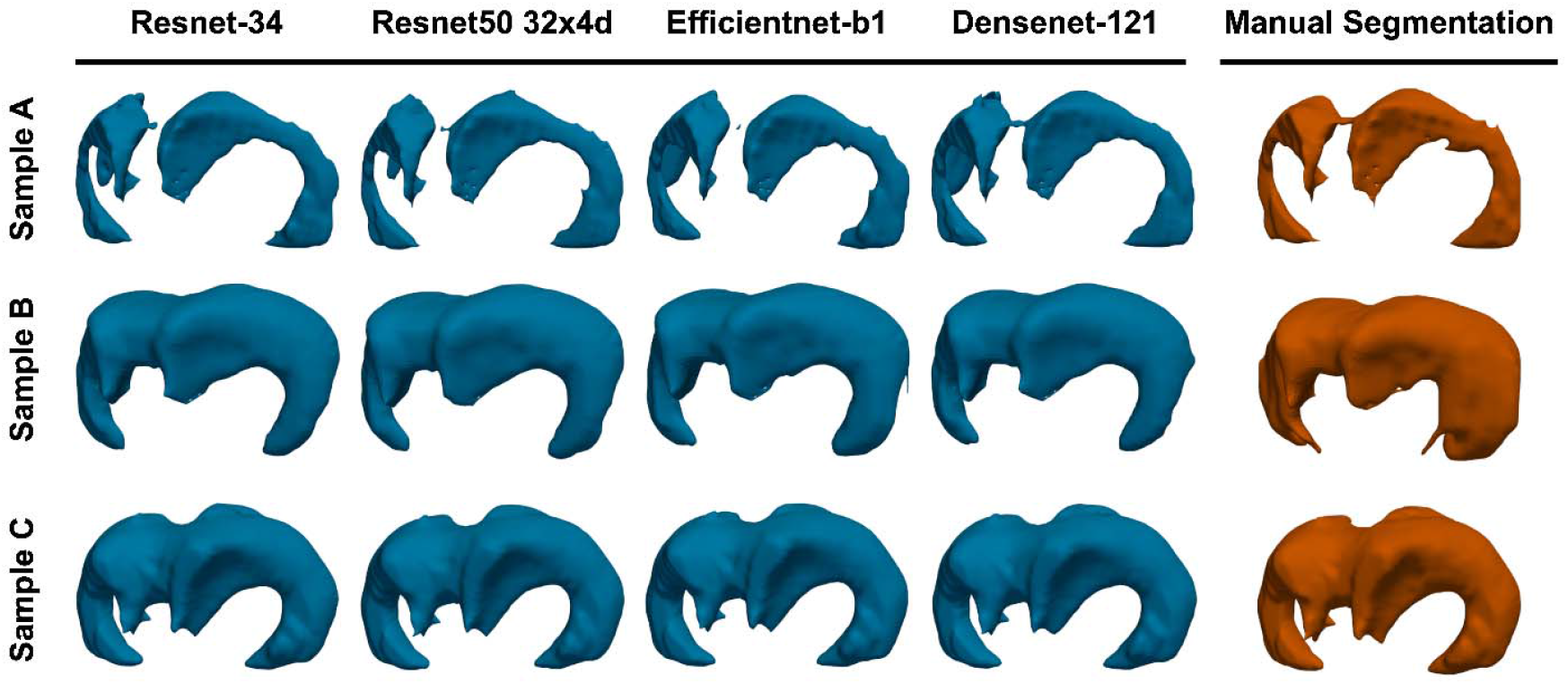
Comparison of 3D reconstructions produced by each U-Net++ variant across three ventricles of varying sizes. Sample A volume = 38.73 mm^3^. Sample B volume = 72.12 mm^3^. Sample C volume = 220.99 mm^3^.

Taken together, the quantitative and qualitative assessments demonstrated no significant differences in performance between the four candidate encoders. However, prior studies have shown that EfficientNet-B1 is both smaller in size and faster in execution compared to the other encoders ^36^. Given these advantages, and with the goal of developing a web-based platform where reduced computational load would improve performance, we selected EfficientNet-B1 as the optimal encoder for subsequent validation analyses.

### Accuracy of Morphological Quantification

Evaluation of the U-Net++ model with the EfficientNet-B1 encoder revealed strong agreement between automated and manual measurements of ventricular morphology. Across all five metrics (volume, normalized volume, surface area, symmetry, and convexity), predicted values closely tracked with the corresponding ground truth measurements. Correlation assessment confirmed this high concordance, with all metrics achieving interclass and Pearson correlation coefficients greater than 0.96 (Table 2). Bland-Altman analysis found that limits of agreement were consistently narrow across all morphological features. No systemic bias was observed, as the mean difference between automated and manual quantifications centered around zero for all metrics. Performance was comparable between rat and mouse samples, although surface area and volume exhibited wider limits of agreement in rats compared to mice (Figure 5).

**Table 2:** Accuracy of Automated Quantifications Compared to Manual Quantifications.

|  | ICC | MAE | Pearson-CC |
| --- | --- | --- | --- |
| **Rats** |  |  |  |
| Ventricle Volume | 0.999 | 4.583 | 0.999 |
| Normalized Ventricle Volume | 0.999 | 0.003 | 0.999 |
| Convexity | 0.990 | 0.018 | 0.992 |
| Surface Area | 0.996 | 13.027 | 0.997 |
| Symmetry | 0.967 | 0.042 | 0.968 |
| **Mice** |  |  |  |
| Ventricle Volume | 0.994 | 0.395 | 0.994 |
| Normalized Ventricle Volume | 0.964 | 0.014 | 0.980 |
| Convexity | 0.983 | 0.009 | 0.983 |
| Surface Area | 0.991 | 2.154 | 0.996 |
| Symmetry | 0.966 | 0.038 | 0.969 |
| **Overall** |  |  |  |
| Ventricle Volume | 0.999 | 3.917 | 0.999 |
| Normalized Ventricle Volume | 0.999 | 0.006 | 0.997 |
| Convexity | 0.990 | 0.017 | 0.991 |
| Surface Area | 0.996 | 11.297 | 0.997 |
| Symmetry | 0.967 | 0.041 | 0.968 |

ICC = interclass correlation coefficient. MAE = Mean Absolute Error, Pearson-CC = Pearson correlation coefficient.

**Figure 5.**
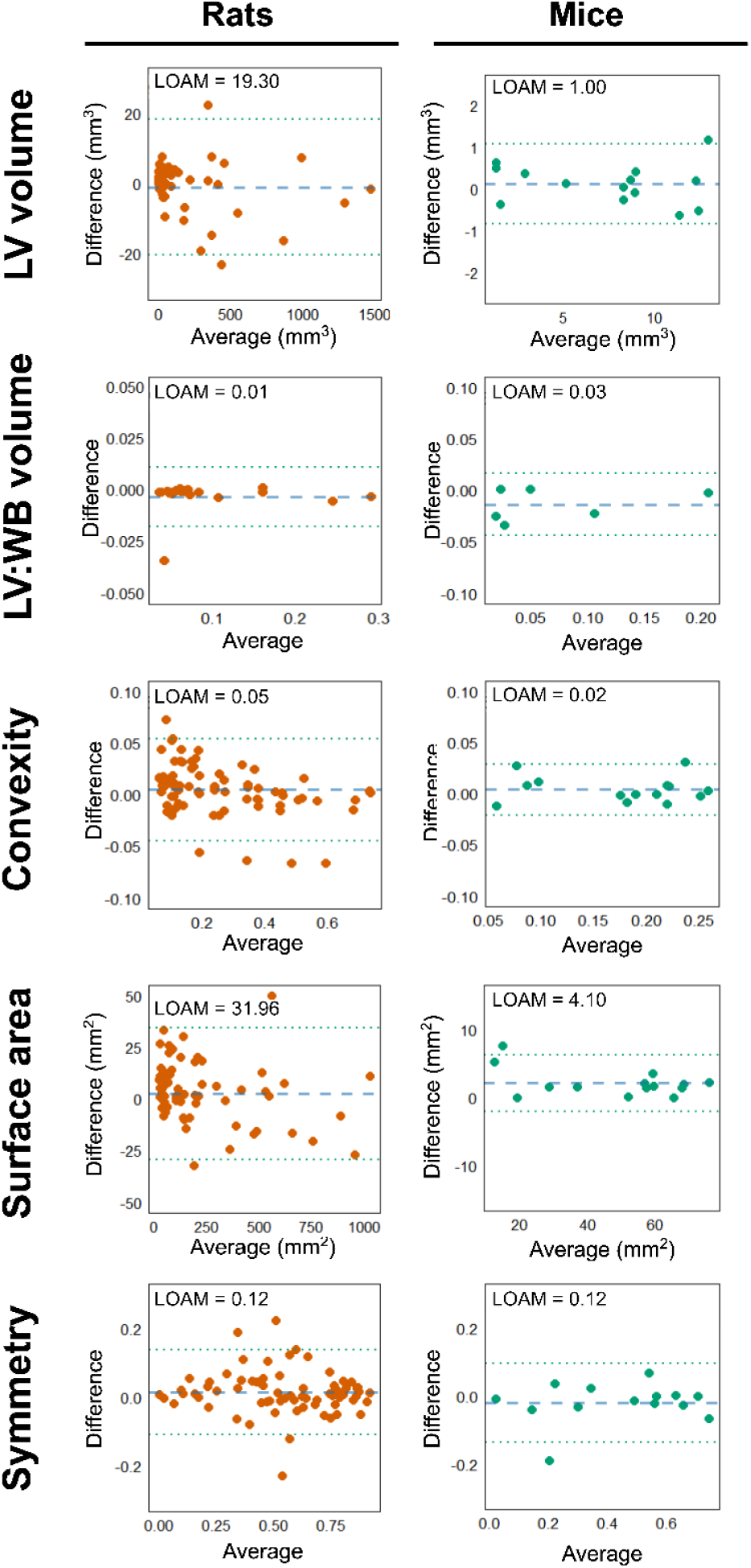
Bland-Altman plots comparing ventricle morphology metrics as calculated by manual segmentation and automated segmentation Plots include average bias (blue dashed line) and 95% limits of agreement (green dashed lines). LOAM = limits of agreement multiplier = 1.96 × SD of the differences.

## DISCUSSION

This study presents a broadly accessible, fully automated deep learning pipeline for the segmentation and morphological quantification of lateral ventricles in rodent models of hydrocephalus. Using a U-Net++ architecture with an EfficientNet-B1 encoder, we developed and validated a model that accurately segments lateral ventricles in MRIs while also creating 3D reconstructions and extracting important morphological metrics. The model demonstrated excellent agreement with manually obtained quantifications, as evidenced by high correlation coefficients, low mean absolute errors, and narrow limits of agreement. This accuracy was maintained across a wide range of ventricle sizes, highlighting the pipeline’s ability to correctly interpret anatomic variability (Supp. Figure 2). To enhance usability, the complete analysis pipeline was integrated into a publicly available website (https://ava-tar.org/).

To the best of our knowledge, this is the first CNN-based tool developed for dedicated lateral ventricle segmentation in preclinical hydrocephalus models. Prior work into automated ventriculomegaly assessment has focused on human MRIs, with tools such as SynthSeg enabling automated segmentation of all intracranial structures ^41^. Similarly, Vahedifard et al. previously developed a CNN-based approach for ventriculomegaly detection in fetal MRIs ^42^. Using these studies as conceptual foundations, our approach extends these concepts into preclinical research and incorporates a more comprehensive quantification pipeline. Along with simple segmentation, our model outputs five metrics - volume, normalized volume, surface area, symmetry, and convexity - each of which is integral in the assessment of hydrocephalus pathology and progression. The strong agreement with manual quantifications across these metrics underscores the accuracy of the model.

A key advantage of our model is its generalizability across rat and mouse MRIs. Although they share general anatomical similarities, species-specific differences in brain size, structure, and image contrast often limit the cross-applicability of digital tools. By explicitly training our model on a large, diverse dataset that included both species and spanned a wide range of ventricular sizes, we achieve a system that is applicable across all rodent hydrocephalus modes. This generalizability expands the utility of the tool, ensuring consistency in analysis across experimental models. In addition, incorporation of total intracranial volumes allows for comparison between different age groups and sizes of animals.

Along with its technical performance, the deployment of our pipeline as an interactive website significantly improves accessibility and usability. Most existing deep learning segmentation models, including SynthSeg, require complex setups, extensive computational knowledge, and access to advanced computing resources.^41^ These requirements can be prohibitive for researchers who lack these skills and resources. In contrast, our web-based implementation allows researchers to easily upload files, execute segmentation and quantification, and visualize 3D reconstructions, all with a common laptop CPU. The website also contains a batch processing feature, making it well-suited for analyzing time-series MRIs used to monitor hydrocephalus progression over time. This streamlined interface minimizes technical barriers, reduces computational costs, and enables standardized analysis.

Our findings suggest that the model performs with slightly higher accuracy on MRIs with larger ventricles compared to those with smaller ones. This likely reflects the greater ease with which the model can identify larger, more clearly defined structures. Additionally, small segmentation inaccuracies have a proportionally greater effect on the overall morphology of small ventricles. An error of just a few pixels can significantly alter the calculated volume in small structures, whereas the same error would be negligible in larger ventricles. The rat MRI dataset encompassed a wider range of ventricular volumes, including many cases with severe ventriculomegaly, whereas the mouse dataset consisted predominantly of smaller ventricles. Consequently, the model’s quantification error was modestly higher in mice across all metrics.

This study has several limitations. First, although the model demonstrated strong generalizability across species, it was trained exclusively on MRIs acquired using a single scanner and protocol. Performance on images from other acquisition settings remains to be validated. Second, the training dataset, while diverse in disease severity, was relatively limited in mouse sample size, which may have constrained model performance in this subgroup. Third, the current model focuses specifically on lateral ventricles and intracranial space. Future studies should extend this framework to include additional intracranial structures or pathologic features that could further enhance its applicability for studying hydrocephalus.

## ACKNOWLEDGEMENTS

This work was supported by the Rudi Schulte Research Institute (J.M.S) and the NIH R01NS110793 (J.M.S.) and 2R01HL128370-09A1 (M.R.M.). The MRI studies presented in this work were carried out using the Small Animal Magnetic Resonance Facility of the Washington University Mallinckrodt Institute of Radiology (S10OD026913).

## AUTHOR CONTRIBUTIONS

S.C. designed the study, organized the raw data, developed all code, analyzed and interpreted data, and wrote the manuscript. S.P. designed the study, performed data collection, analyzed and interpreted data, and wrote the manuscript. O.L., M.P., G.L.H, and A.Z. performed data collection. M.R.M, J.D.Q, and A.N. supervised the project and provided expert guidance. J.M.S. designed the study, supervised the project, and provided expert guidance.

## DATA AVAILABILITY

The images used for model development and testing are available from the corresponding author upon reasonable request. The source code used to develop the automated alignment assessment web app is publicly available at https://github.com/sundeep3257/AVA-TAR. This includes the fully trained model used for ventricle segmentation.

## ABBREVIATIONS

AVA-TAR: Automated Ventricular Analysis Tool for Rodents
CNN: Convolutional Neural Network
CSF: Cerebrospinal Fluid
FOV: Field of View
GPU: Graphics Processing Unit
HB: Hemoglobin
ICV: Intracranial Volume
IoU: Intersection over Union
IVH: Intraventricular Hemorrhage
LV: Lateral Ventricle
MAE: Mean Absolute Error
MRI: Magnetic Resonance Imaging
PHH: Post-Hemorrhagic Hydrocephalus
SD: Standard Deviation
U-Net++: Nested U-Net Architecture

## SUPPLEMENTARY FILES

Table 1. Parameter settings for used model development.

Figure 1. Training and validation loss over successive epochs of each U-net model during training.

Figure 2. Distribution of ventricle volumes used for training models.

