## Supplemental Figure 1 for "Automated Ventricle Assessment via Three-dimensional Anatomical Reconstruction (AVA-TAR): a computational toolkit for autonomous lateral ventricle assessment in preclinical hydrocephalus models"

### Slide 1
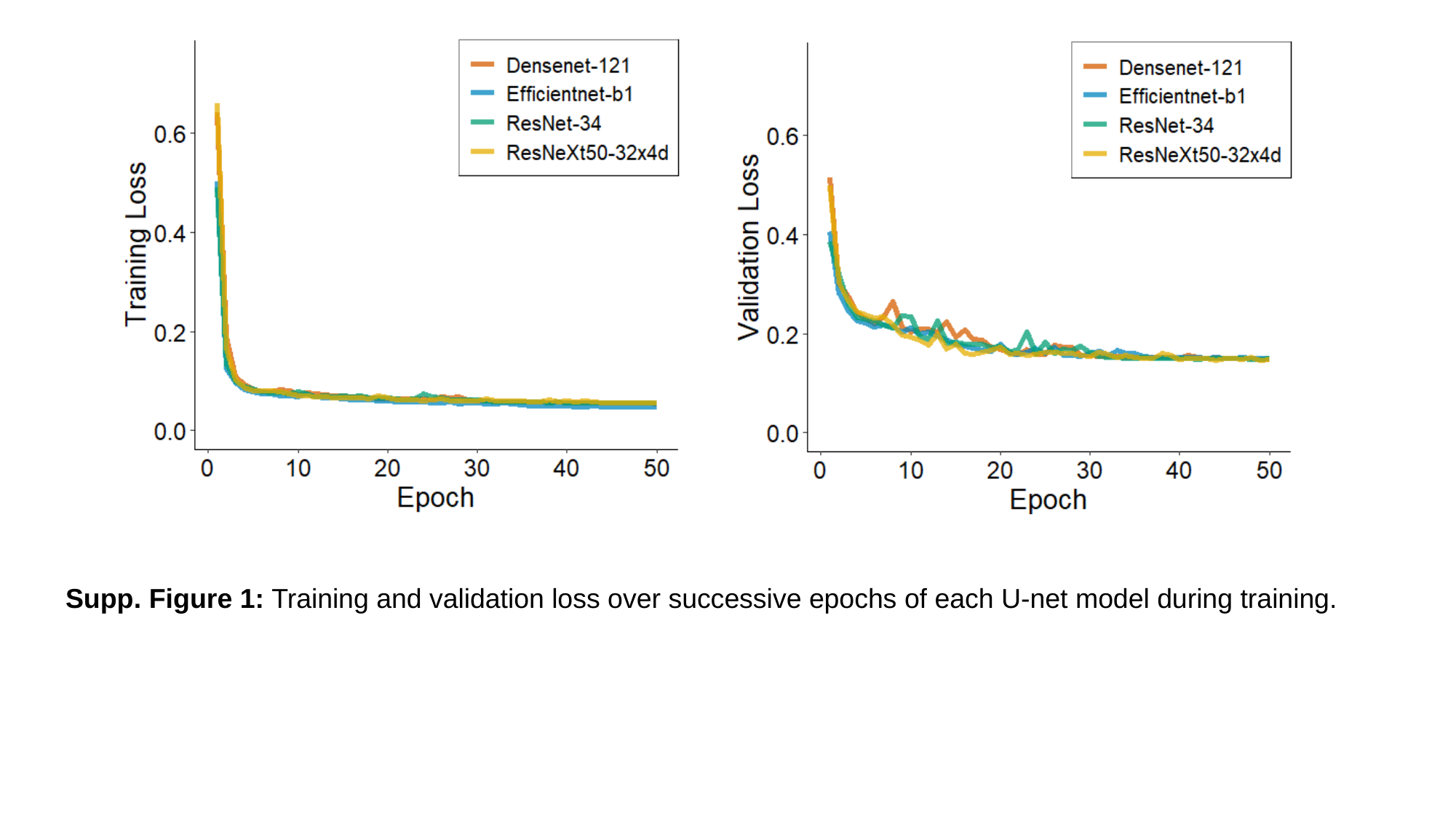

Supp. Figure 1: Training and validation loss over successive epochs of each U-net model during training.

### Slide 2
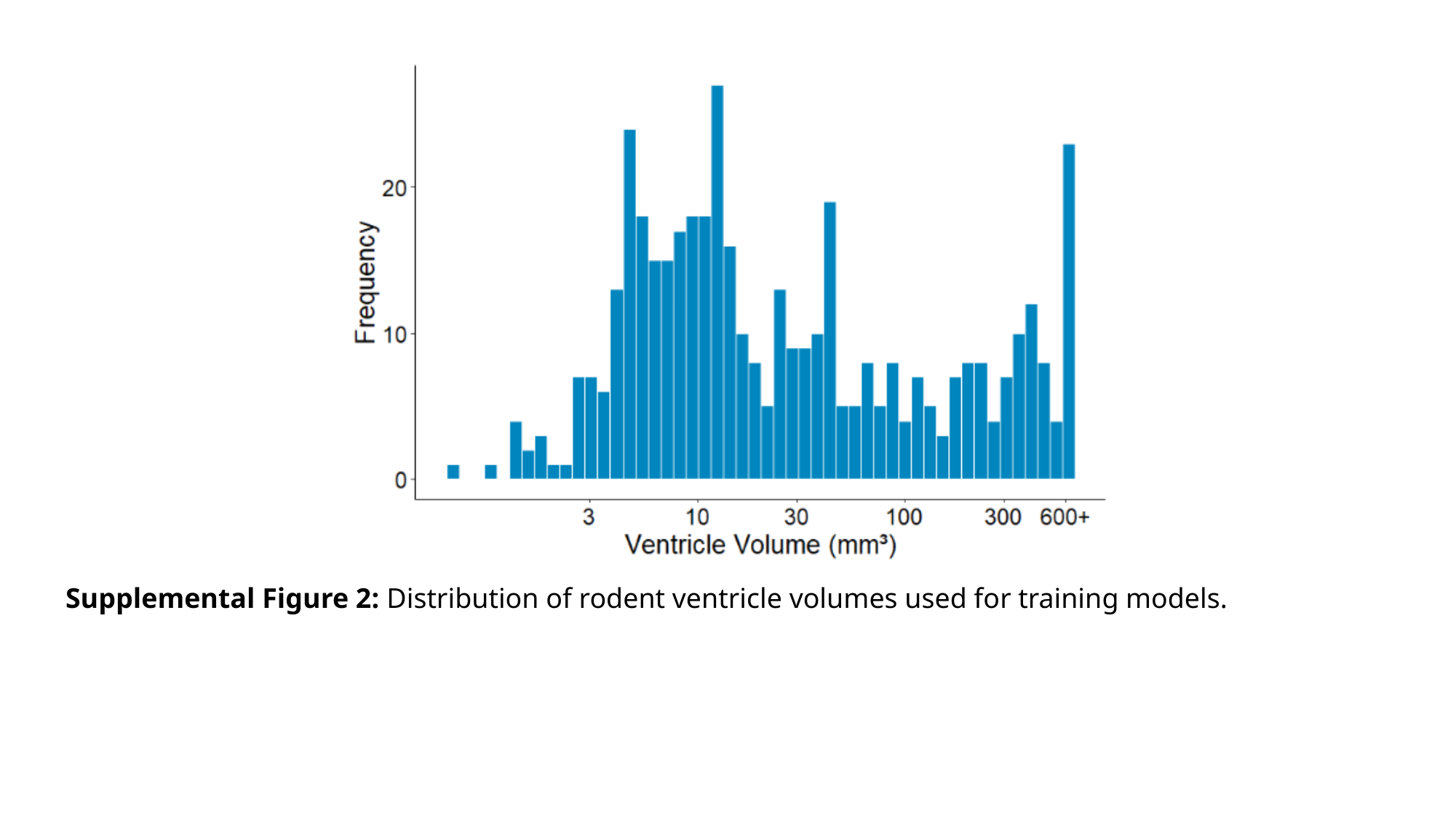

Supplemental Figure 2: Distribution of rodent ventricle volumes used for training models.
