## Supplemental Table 1 for "Automated Ventricle Assessment via Three-dimensional Anatomical Reconstruction (AVA-TAR): a computational toolkit for autonomous lateral ventricle assessment in preclinical hydrocephalus models"

**Supplemental Table 1:** Parameter Settings for Model Development

| **Parameter** | **Value Selected** | **Parameter Description** |
| --- | --- | --- |
| **Training Setup** | | |
| optimizer | optim.Adam() | Uses the Adam optimizer for adaptive learning rate updates |
| lr | 0.0001 | Sets a small learning rate to ensure gradual and stable learning |
| batch_size | 4 | Small batch size chosen to fit high-resolution MRI slices into GPU memory |
| num_epochs | 20 | Model is trained for 20 epochs |
| **Model Architecture** | | |
| model | smp.UnetPlusPlus() | Employs the U-Net++ architecture |
| encoder_name | resnet34  resnet50_32x4d  efficientnet-b1  densenet121 | Specifies the backbone for feature extraction in the U-Net++ architecture |
| encoder_weights | imagenet | Initializes the encoder with pre-trained ImageNet weights |
| **Loss Function** | | |
| BCEWithLogitsLoss | | Binary Cross Entropy with logits is used for stable probabilistic output in binary segmentation |
| DiceLoss(mode='binary') | | Measures overlap between predicted and true masks |
| JaccardLoss(mode='binary') | | Penalizes differences in shape and size between predicted and ground truth masks (also known as IoU loss) |
| loss = 0.5(BCE) + 1.0(Dice) + 1.0(IoU) | | A custom loss function that balances BCE (0.5), Dice (1.0), and IoU (1.0) to optimize both pixel-wise and spatial accuracy |

BCE: binary cross entropy. IoU: intersection over union
